# Chronic *In Vivo* Imaging of a Focal Cortical Dysplasia Mouse Model Reveals Progressive Neuronal Loss with No Senolytic or Antiseizure Efficacy Following Dasatinib–Quercetin Treatment

**DOI:** 10.64898/2026.05.19.726138

**Authors:** C.V.L. Olson, N. Shariati, N. Procházková, K. Čížek, M. Řehořová, J. Populová, J.T. Rozlivková, S. Wang, B. Ricketts, B. Kučerová, J. Kudláček, B. Straka, P. Jiruška, O. Novák

## Abstract

**Objective:** Focal cortical dysplasia (FCD) type II is a common cause of pediatric drug-resistant epilepsy with no disease-modifying treatment. It arises from somatic mutations in the mTOR pathway that produce a mosaic population of mTOR-hyperactive dysmorphic neurons (DNs) with a senescence phenotype. A recent report showed that oral dasatinib– quercetin (DQ), both putative senolytics, selectively eliminates these neurons and abolishes seizures in an *Mtor*^p.S2215F^ model. However, this has not been tested in a second FCD model and neuron loss was shown only by cross-sectional histology.

**Methods:** Using chronic cranial windows and longitudinal *in vivo* two-photon imaging, we tracked genetically labeled neurons over eight weeks. In a mosaic *MTOR*^p.L2427P^ FCD model on a C57BL/6J background, we combined imaging with continuous video-EEG, immunohistochemistry (IHC), and LC–MS/MS pharmacokinetics in 47 mice. Seizure and pharmacokinetic sampling were extended to a smaller Swiss/CD1 cohort, the strain used previously.

**Results:** Imaging revealed progressive loss of mTOR_mut_ (16.7%) and non-mTOR_mut_ (13.1%) neurons, with minimal loss in unmatched non-FCD controls. In IHC, lesions contained β-galactosidase-positive (B-Gal^+^) cells, but oral DQ showed no evidence of eliminating DNs or reducing seizure frequency, duration, or burden; seizures also persisted in CD1 mice. B-Gal^+^ cells tended to be less dense after DQ, but not significantly after correction. These data are more consistent with a senomorphic than a senolytic effect. Brain dasatinib (median 8.5 nM) exceeded its Src/ABL IC_50_, whereas quercetin (median 12 nM) was nearly two orders of magnitude below its BCL-XL affinity.

**Significance:** Longitudinal tracking of single neurons shows progressive loss of labeled neurons in an FCD model, whereas the reported senolytic and antiseizure effects of DQ were not reproduced. DQ should not be assumed to act as a senolytic across FCD models, strains, or *MTOR* variants. Dependence on strain, mutation, or brain exposure bears on clinical translation.

**Key Points**

- Longitudinal *in vivo* two-photon imaging showed progressive loss of mTOR-mutant neurons in an FCD type II mouse model
- Oral dasatinib–quercetin treatment did not eliminate dysmorphic neurons or reduce seizures in C57BL/6J mice; seizures persisted in CD1 mice
- Dasatinib–quercetin did not eliminate neurons or abolish seizures in an MTOR^p.L2427P^ FCD model, so its efficacy may not generalize

## Introduction

Focal cortical dysplasia type II (FCD II) is the most common histopathological diagnosis in children undergoing epilepsy surgery^1^, and roughly three-quarters of affected children meet the International League Against Epilepsy (ILAE) criteria for drug-resistant epilepsy^2,3^. Surgical resection remains the only potentially curative option, yet only about two-thirds of operated patients with FCD II are free of disabling seizures at five years^4,5^. FCD II arises from somatic mutations in genes of the PI3K–AKT–mTOR pathway, most commonly *MTOR* itself, producing a mosaic population of mTOR-hyperactive dysmorphic neurons (DNs) and, in FCD IIb, balloon cells (BCs), intermingled with histologically normal neurons^6,7^. Targeting the pathway directly has had limited success in FCD II; everolimus is effective in TSC-associated epilepsy^8^, but a prospective trial of sirolimus in FCD II did not meet its primary endpoint^9^. Systemic mTOR suppression alone thus does not fully address the epileptogenic process, motivating the search for mechanistically distinct, disease-modifying treatments.

One such strategy follows from the recognition that mTOR-hyperactive DNs display hallmarks of cellular senescence. Senescence is a stable, largely irreversible cell-cycle arrest entered in response to various stressors that converge on DNA damage. Senescent cells remain metabolically active and adopt a senescence-associated secretory phenotype (SASP), a sustained output of pro-inflammatory cytokines, chemokines, and matrix-remodeling factors capable of altering the surrounding tissue^10,11^. First characterized as tumor-suppressive, senescence is increasingly implicated in chronic, non-neoplastic disease across organ systems, including the central nervous system^10^.

Constitutive mTORC1 activity is well-positioned to drive this state. It imposes proteotoxic stress through excess ribosome biogenesis, suppresses ULK1-mediated autophagy so that ROS-generating mitochondria accumulate and sustain DNA damage signaling and SASP output^12–14^, and engages both principal senescence arrest programs (p53/p21 via 4EBP1^15^; p16/Rb via S6K1^16^). Building on earlier reports of senescence markers in FCD tissue^17^ and a prior study using the same dosing^18^, Ribierre and colleagues showed that DNs and BCs in human FCD IIb specimens are positive for senescence-associated β-galactosidase (B-Gal), p53, and p16^19^. In a mosaic *Mtor*^p.S2215F^ mouse model, a nine-day course of oral dasatinib plus quercetin (DQ) reduced pS6^+^ and B-Gal^+^ density by roughly 70–80%, suppressed six of eight assayed SASP cytokines, and rendered animals seizure-free within one month^19^. Because DQ is already in clinical trials for other indications^20^, these findings raised the prospect of rapid translation to FCD.

However, the evidence for neuronal elimination rests on cross-sectional histology. Whether mTOR-mutant neurons are progressively lost over the course of disease, and whether such loss is druggable, has not been established with a method capable of tracking the same cells over time. We addressed this directly by performing longitudinal *in vivo* two-photon imaging of an FCD mouse model, tracking individual genetically labeled neurons across eight weeks alongside continuous video-EEG monitoring. Although our model carried a different recurrent variant (*MTOR*^p.L2427P^)^21–23^ on a different background (C57BL/6J rather than Swiss/CD1), we used the same platform, together with multiplexed immunohistochemistry (IHC) and targeted liquid chromatography tandem mass spectrometry (LC–MS/MS), to test whether the DQ senolytic effect generalizes to this second, independent FCD model. We additionally obtained continuous video-EEG and LC– MS/MS data from a smaller cohort of *MTOR*^p.L2427P^ mice on the Swiss/CD1 background used by Ribierre *et al.*^19^.

## Materials and Methods

### Animals and FCD model

All experiments were performed under the Animal Care and Animal Protection Law of the Czech Republic, compatible with European Union directive 2010/63/EU, and approved by the Ethics Committees of the Second Faculty of Medicine (Project License No. MSMT-26023/2023-5). Mosaic FCD lesions were induced by *in utero* electroporation (IUE) at E14.5 of CAG-mTOR^p.L2427P^-IRES-CreERT2 and CAG-EGFP plasmids into C57BL/6J × LSL-tdTomato (Ai9) embryos; tamoxifen from week 5 activated CreERT2, restricting tdTomato expression to mTOR-mutant neurons, while EGFP labeled all electroporated cells. Swiss/CD1 (CD-1/ICR) embryos received the same plasmid mixture for the EEG and LC–MS/MS cohorts and are referred to as CD1 throughout. Non-FCD controls were VIP-Cre × Ai9 mice injected with AAV-hSyn-jGCaMP7f.

Animals used in each analysis are listed in Table 1.

**Table 1.**
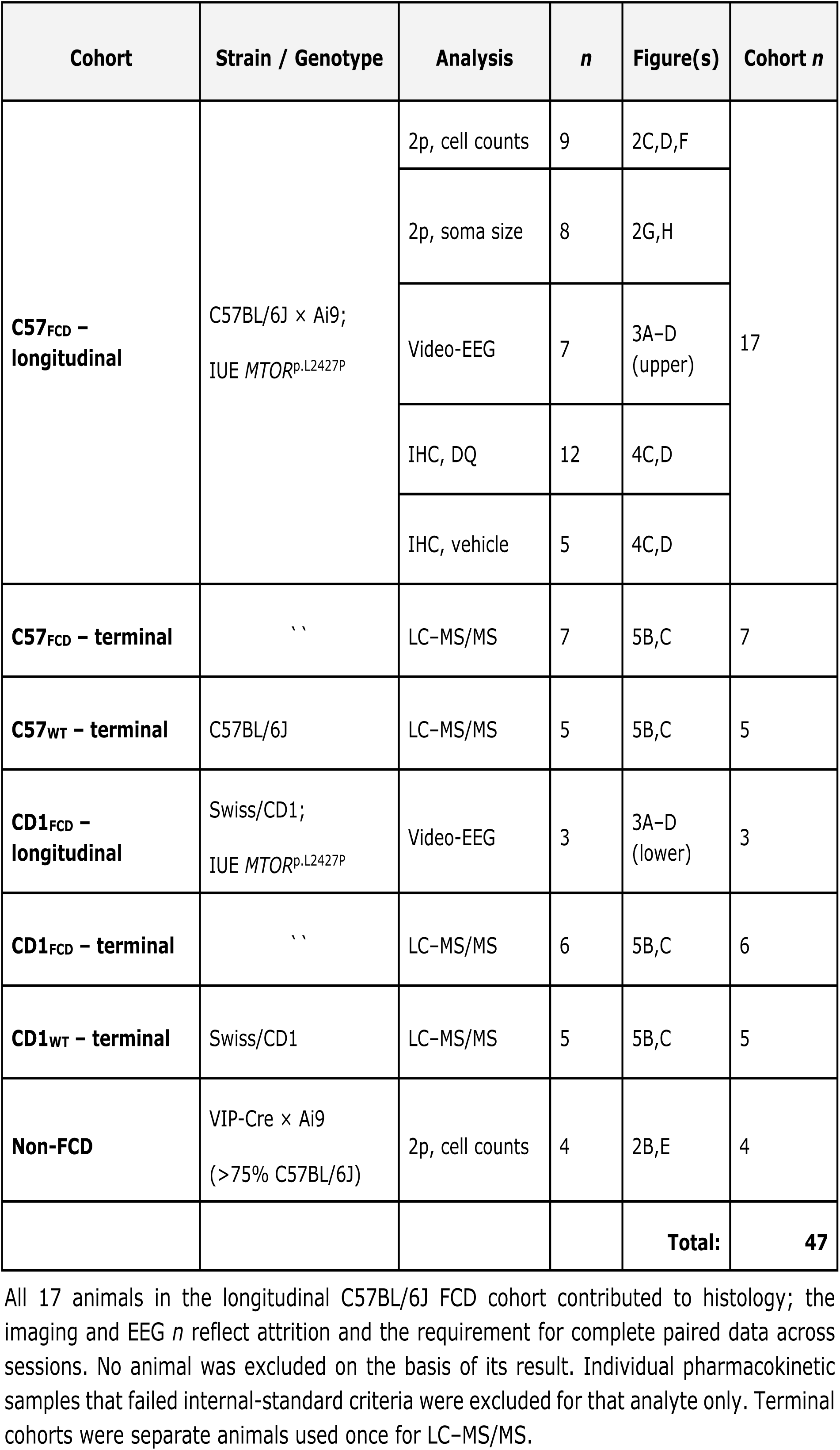
Animals used in each analysis.

| Cohort | Strain / Genotype | Analysis | *n* | Figure(s) | Cohort *n* |
| --- | --- | --- | --- | --- | --- |
| **C57<sub>FCD</sub> – longitudinal** | C57BL/6J × Ai9;<br>IUE *MTOR*<sup>p.L2427P</sup> | 2p, cell counts | 9 | 2C,D,F | 17 |
|  |  | 2p, soma size | 8 | 2G,H |  |
|  |  | Video-EEG | 7 | 3A–D (upper) |  |
|  |  | IHC, DQ | 12 | 4C,D |  |
|  |  | IHC, vehicle | 5 | 4C,D |  |
| **C57<sub>FCD</sub> – terminal** | `` | LC–MS/MS | 7 | 5B,C | 7 |
| **C57<sub>WT</sub> – terminal** | C57BL/6J | LC–MS/MS | 5 | 5B,C | 5 |
| **CD1<sub>FCD</sub> – longitudinal** | Swiss/CD1;<br>IUE *MTOR*<sup>p.L2427P</sup> | Video-EEG | 3 | 3A–D (lower) | 3 |
| **CD1<sub>FCD</sub> – terminal** | `` | LC–MS/MS | 6 | 5B,C | 6 |
| **CD1<sub>WT</sub> – terminal** | Swiss/CD1 | LC–MS/MS | 5 | 5B,C | 5 |
| **Non-FCD** | VIP-Cre × Ai9<br>(>75% C57BL/6J) | 2p, cell counts | 4 | 2B,E | 4 |
|  |  |  |  | **Total:** | **47** |

All 17 animals in the longitudinal C57BL/6J FCD cohort contributed to histology; the imaging and EEG *n* reflect attrition and the requirement for complete paired data across sessions. No animal was excluded on the basis of its result. Individual pharmacokinetic samples that failed internal-standard criteria were excluded for that analyte only. Terminal cohorts were separate animals used once for LC–MS/MS.

### Treatment

Dasatinib (12 mg/kg) and quercetin (50 mg/kg) in 30% PEG300 / 5% Tween-20, or vehicle, were given by oral gavage for four consecutive days, a two-day intermission, and a further five consecutive days—nine dosing days in total, matching the dose and regimen reported by Ribierre *et al.*^19^. Vehicle was administered during weeks 16–18 and DQ during weeks 20–22, so each animal served as its own control.

### *In vivo* imaging and EEG

A chronic cranial window and EEG electrodes were implanted at approximately week 12. Two-photon Z-stacks were acquired on a custom high-resolution microscope in six sessions between weeks 16 and 24. Stacks were co-registered across sessions with ITK-elastix and cells tracked manually in FIJI. Continuous video-EEG (0.1 Hz–1.6 kHz, 5 kHz sampling) was recorded from week 14 to week 24 and scored manually, blind to the recording phase. Electrographic seizures were defined as rhythmic epileptiform discharges of at least 2–3× baseline amplitude, >2 Hz, and ≥10 s, with evolution in amplitude and frequency, recorded at lesional and contralateral electrodes, and confirmed on synchronized video; behavioral semiology followed a modified Racine scale^24,25^.

### Histology

After the final imaging session, about two weeks after the last DQ dose, brains were perfused, sectioned, and processed for B-Gal, detected as X-Gal crystalline fluorescence in the far-red channel^26^, and for pS6 and NeuN by immunohistochemistry. Spinning-disk confocal maximum intensity projections were segmented with Cellpose 3.1.1.2 and densities computed within the EGFP-defined lesion.

### Pharmacokinetics

Dasatinib and quercetin were quantified in brain and serum from 23 mice by LC–MS/MS with deuterated internal standards on a Triple Quad 6500+ system, after the fourth of four daily doses, with calibration matrix from undosed animals. Samples lacking internal-standard signal were excluded for that analyte only.

### Statistics

Analyses were performed in MATLAB R2024b, Python 3.11.5, and GraphPad Prism 10.6.1, with histological analyses blind to treatment group, using two-sided nonparametric tests given the small group sizes. Values are median (IQR) or mean±SEM. LC–MS/MS concentrations were compared across strain and genotype with stratified rank (van Elteren-type) permutation tests. Bonferroni correction was applied across histological markers; *P*<0.05 was considered significant. No *a priori* sample-size calculation was performed. Sensitivity was assessed *post hoc* as the minimum detectable effect of each paired within-animal comparison (two-sided α=0.05, 80% power, paired-*t* approximation using the observed SD of the differences).

Full procedural details are provided in the supplementary *Extended Methods*.

## Results

The experimental design is summarized in Figure 1 and described in *Materials and Methods*. Tamoxifen-induced Cre recombination marks mTOR-mutant neurons with tdTomato, whereas the electroporated cells without the mutant plasmid express only EGFP. Throughout, EGFP⁺ denotes EGFP⁺/tdTomato⁻ cells; tdTomato^+^ cells can contain EGFP. Each animal received vehicle (weeks 16–18) and then DQ (weeks 20–22), as in Ribierre *et al.*^19^, so the vehicle phase served as a within-animal control; vehicle-only controls received vehicle in both phases. Pharmacokinetics were measured in separate animals (Figure 5; Table 1).

**Figure 1.**
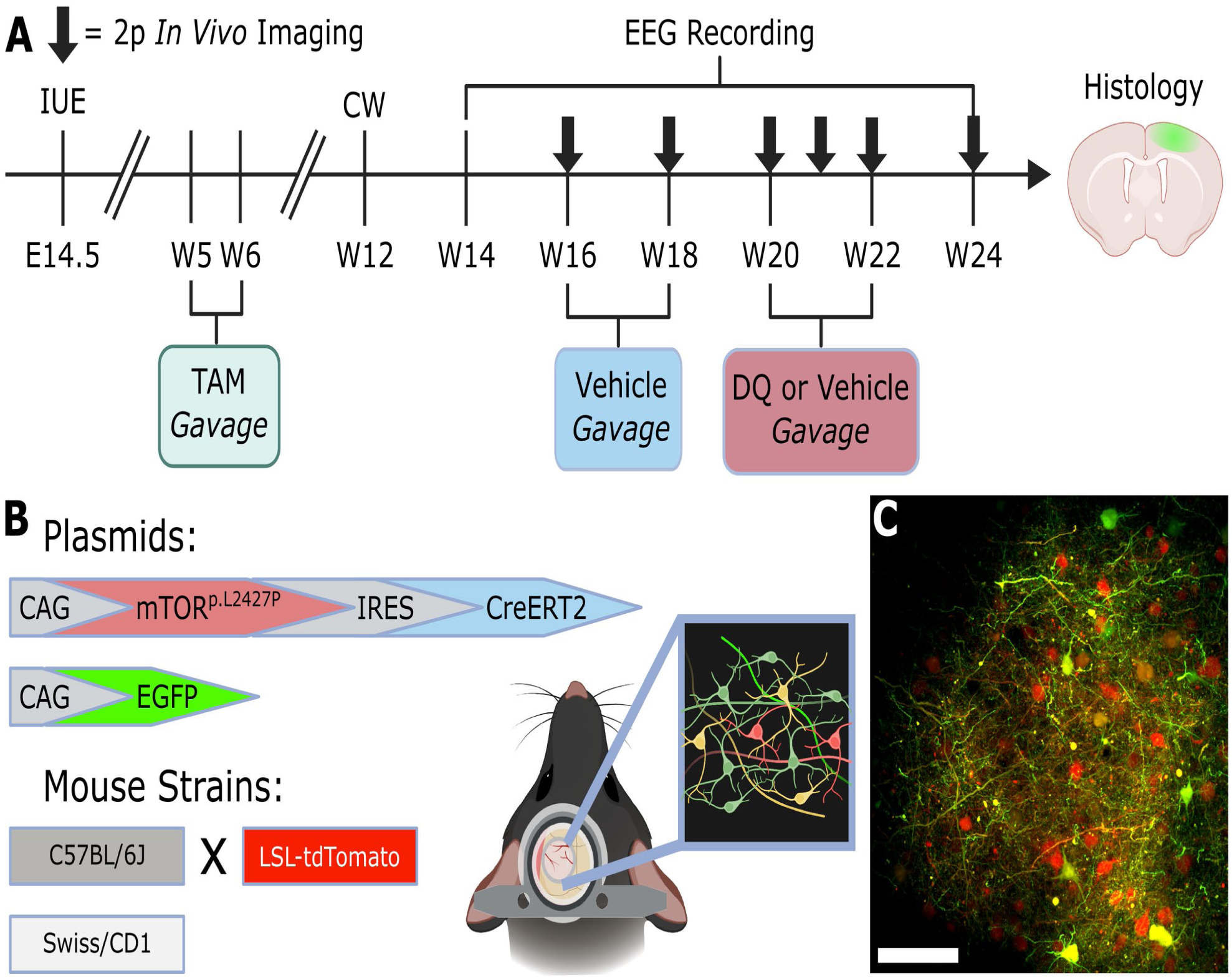
Experimental design and models. **A**) Timeline of the longitudinal experiment in C57BL/6J mice (axis breaks denote compressed intervals; not to scale). FCD was induced by IUE at embryonic day 14.5 (E14.5). Tamoxifen (TAM) was given by oral gavage starting at week 5 to activate CreERT2. A cranial window (CW) and EEG electrodes were implanted at approximately week 12. Continuous video-EEG was recorded from week 14 to week 24 (bracket), and two-photon (2p) *in vivo* imaging sessions (arrows) were acquired at weeks 16, 18, 20, 21, 22, and 24. Oral gavage of vehicle (blue) at weeks 16–18 was followed by DQ or vehicle again (red; vehicle in vehicle-only controls) at weeks 20–22. Brains were collected after week 24 for histology. **B**) Electroporated plasmids (CAG-mTOR^p.L2427P^-IRES-CreERT2 and CAG-EGFP) and host strains: C57BL/6J crossed with a Cre-dependent LSL-tdTomato Ai9 reporter line (imaging, EEG, histology, and LC–MS/MS) and Swiss/CD1 (EEG and LC–MS/MS only). EGFP labels all electroporated neurons; tdTomato is expressed only after Cre recombination and therefore marks mTOR-mutant neurons. Schematic depicts the chronic cranial window used for 2p imaging and a representation of the visible field of view (FOV) under 2p imaging. **C**) Example *in vivo* 2p FOV in an FCD mouse (EGFP, green; tdTomato, red); scale bar = 100 μm. Subplots **A** and **B** were created with https://BioRender.com.

### Oral DQ Did Not Eliminate DNs in a C57BL/6J FCD Type II Model

To assess whether DQ induces selective elimination of DNs, we performed longitudinal *in vivo* two-photon imaging of genetically labeled cells at intervals of one to two weeks between weeks 16 and 24 (Figure 2A).

**Figure 2.**
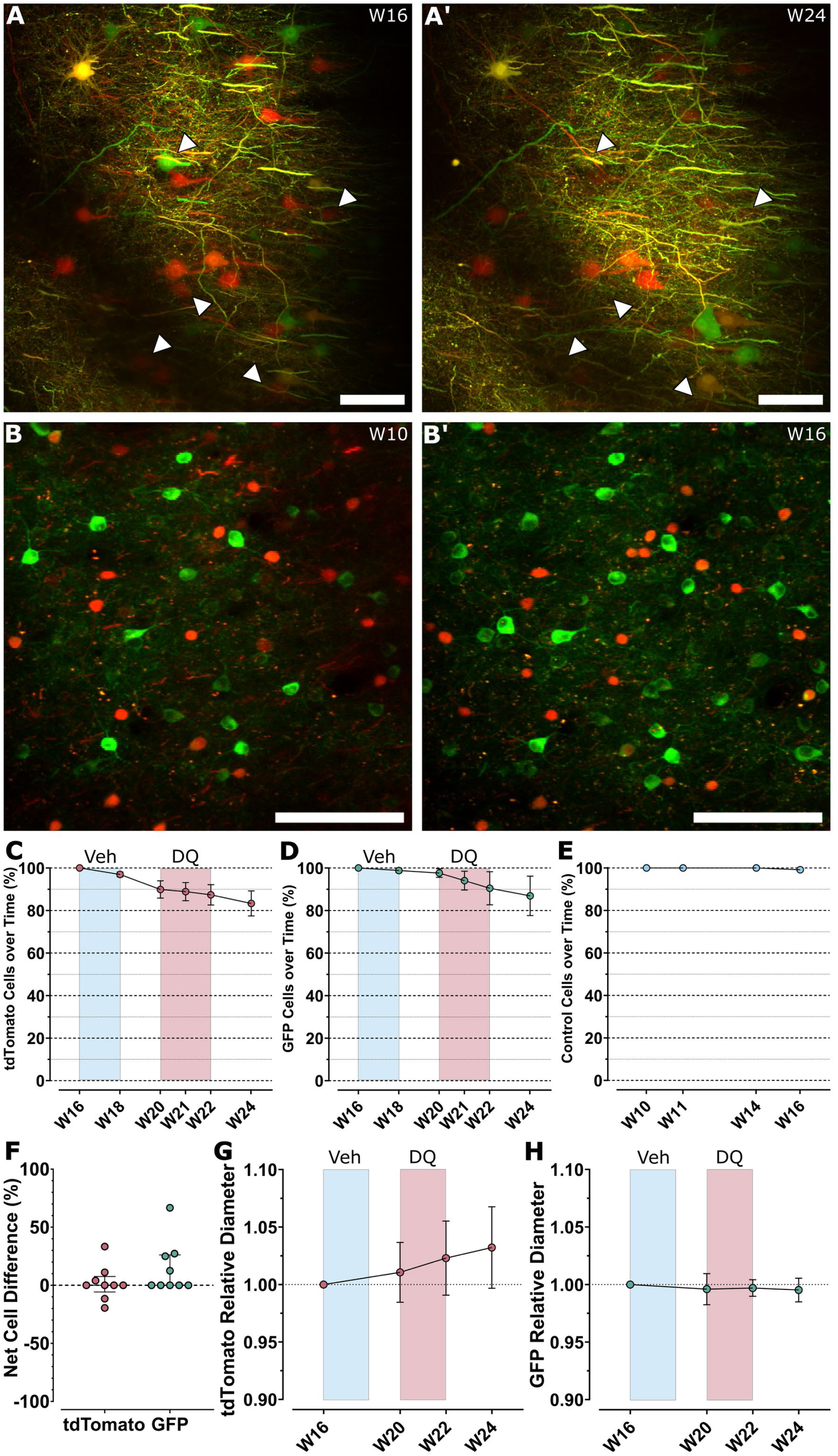
Longitudinal *in vivo* two-photon imaging of electroporated and mTOR-mutant neurons in the C57BL/6J FCD model. **A**, **A’**) Maximum intensity projections (MIPs) of the same FOV in an FCD mouse at week 16 (**A**) and week 24 (**A’**); EGFP (green), tdTomato (red); arrows denote cells that were lost between weeks 16–24. **B**, **B’**) MIPs from a non-FCD control mouse at weeks 10 (**B**) and 16 (**B’**); neurons expressing the calcium indicator GCaMP7f (green) and VIP^+^ interneurons (tdTomato, red); controls come from a separate experiment. **A**–**B’**) Scale bars = 100 µm. **C**–**E**) Cells remaining as a percentage of the cells counted in the first session for tdTomato^+^ (**C**) and EGFP^+^ (**D**) neurons in FCD mice (n = 9) and for GCaMP7f^+^ neurons in control mice (**E**; n=4); points and whiskers denote means ± SEM, weighted by baseline cell number per animal; controls in (**E**) were imaged at weeks 10, 11, 14, and 16 and received no gavages; at week 24, tdTomato⁺ neurons had declined by 16.7% (95% CI 3.6–31.1) and EGFP⁺ neurons by 13.1% (95% CI 1.5– 39.6; cluster bootstrap over animals, n = 9). **F**) Net difference in cell loss between the DQ and vehicle phases, |loss|DQ − |loss|Vehicle, for tdTomato^+^ and EGFP^+^ neurons; each point is one animal (n = 9); lines and error bars denote the median and IQR; Hodges–Lehmann median differences (percentage points, exact 96.1% CI): tdTomato⁺ 0.0 (−9.8 to 16.7); EGFP⁺ 12.5 (0.0 to 33.3). **G**, **H**) Soma diameters relative to week 16 for tdTomato^+^ (**G**) and EGFP^+^ (**H**) neurons across four imaging sessions (ntdTomato = 8 animals, nEGFP = 6 animals); points and whiskers denote weighted means±SEM. In **C**–**E**, **G**, and **H**, blue and red shading denote the vehicle and DQ phases, respectively.

Imaging revealed a progressive reduction in both tdTomato^+^ (16.7%) and EGFP^+^ (13.1%) neuron counts over the eight-week experimental period (Figure 2C,D). This decrease was statistically significant in both populations (tdTomato: Friedman *Q*=23.10, *P*<0.001; EGFP: Friedman *Q*=17.93, *P*=0.003), in line with the reduced neuronal density reported in FCD lesions^23^. That reduction has been attributed to tissue expansion rather than cell death, so density alone cannot indicate neuronal loss^27^. Because the Cre-dependent tdTomato and the constitutively expressed EGFP populations declined in parallel, the loss is unlikely to reflect silencing of a single reporter, although fluorescence data alone cannot exclude a general decline in transgene expression. To assess whether this reflects disease state rather than imaging drift of transgene expression, we compared against a small cohort of non-FCD controls (AAV-hSyn-jGCaMP7f into VIP-Ai9 mice; Figure 2B), which showed stable counts over a similar duration (Kruskal–Wallis *H*=1.094, *P*=0.86; Figure 2E).

The difference between DQ-phase and vehicle-phase cell loss within animals showed no DQ-attributable effect (Figure 2F). Within-animal analyses showed no detectable DQ effect on either cell population (tdTomato: Wilcoxon *W*=1.000, *P*>0.99; EGFP: Wilcoxon *W*=10.00, *P*=0.12). For EGFP, losses were equal in the two phases in five of nine animals and larger during DQ in four. Relative soma diameters showed no treatment-associated changes in either population (tdTomato: Friedman *Q*=6.6, *P*=0.09; EGFP: Friedman *Q*=1.4, *P*=0.77; Figure 2G,H); tdTomato^+^ diameter trended upward by about 3% by week 24 (1.03±0.04 relative to week 16), whereas EGFP^+^ diameter was stable (1.00±0.01). Together, these longitudinal *in vivo* data show that oral DQ does not induce detectable neuronal death or atrophy in this FCD model.

### Oral DQ Did Not Reduce Seizure Burden in a C57BL/6J FCD Type II Model

We next evaluated whether oral DQ affects seizure frequency or burden. We recorded continuous video-EEG throughout baseline, vehicle, inter-treatment, treatment, and post-treatment phases (average recorded EEG hours per C57 animal: 1360±70), enabling within-animal longitudinal analysis with minimal sampling bias.

Seizure frequency did not differ for DQ-treated animals at any phase of the experiment (Friedman *Q*=2.057, *P*=0.73; Figure 3A upper), nor did mean seizure duration (Friedman *Q*=5.143, *P*=0.27; Figure 3B upper), or seizure burden, given as seizing minutes per 24 hours (Friedman *Q*=1.829, *P*=0.77; Figure 3C upper). Comparing DQ-phase and vehicle-phase seizure frequencies within the same animals likewise revealed no DQ-attributable reduction (Wilcoxon *W*=−12.00, *P*=0.38; Figure 3D upper).

**Figure 3.**
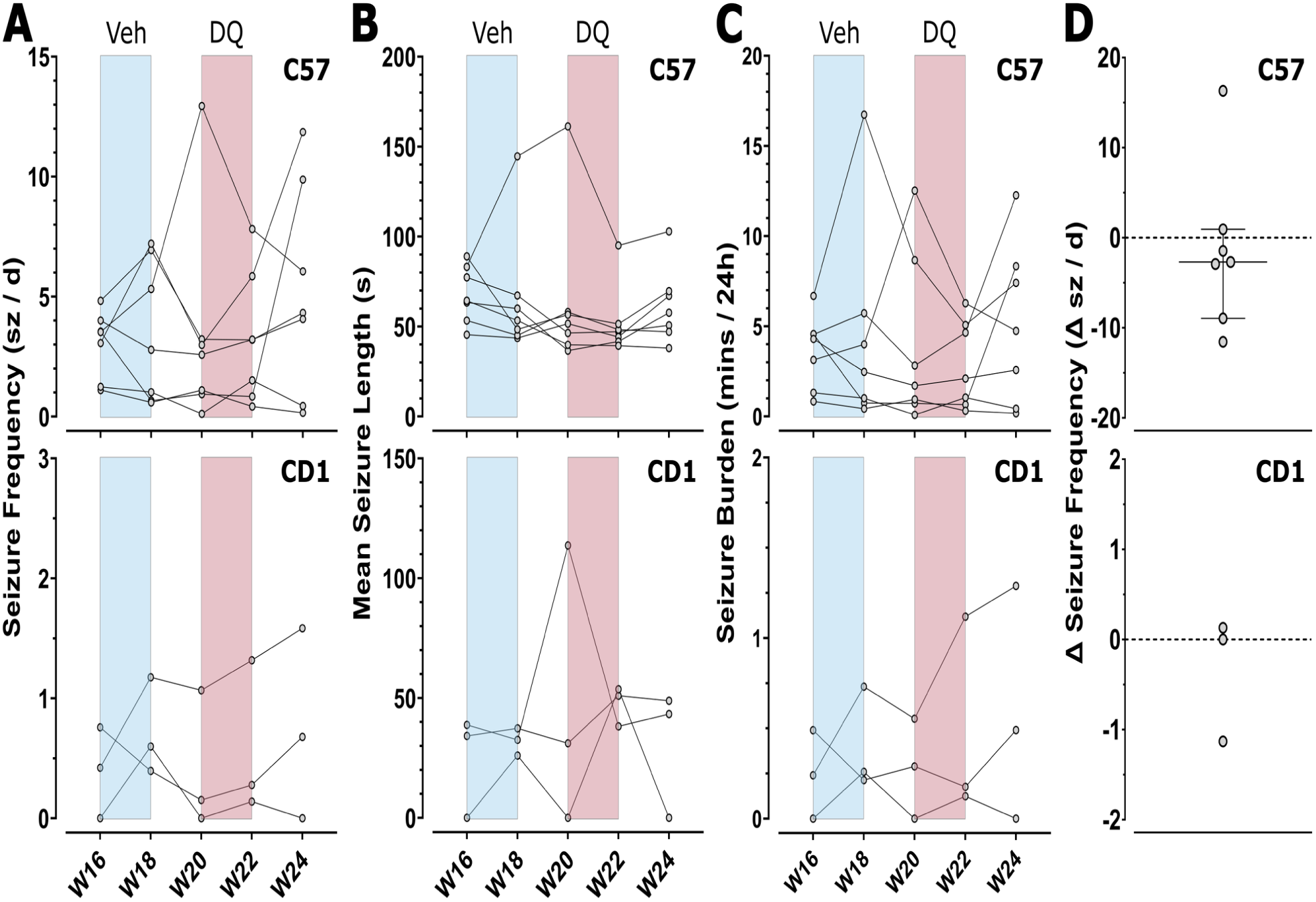
Continuous video-EEG seizure monitoring across vehicle and DQ phases in C57BL/6J and Swiss/CD1 FCD mice. **A**–**C**) Seizure frequency (**A**), mean seizure duration (**B**), and seizure burden (**C**; seizing minutes per 24 h) for C57BL/6J (top; n = 7) and Swiss/CD1 (bottom; n=3) mice; each line is one animal and each point is the per-animal value for the indicated preceding two weeks; weeks without seizures are plotted as zero in **A**– **C**; blue and red shading denote the vehicle and DQ phases, respectively. **D**) Within-animal change in seizure frequency during DQ relative to vehicle, Δ = |seizures/day|DQ − |seizures/day|Vehicle; each point is one animal; lines and error bars denote the median and IQR (C57; Hodges–Lehmann median difference −2.4 seizures/day, exact 95.3% CI −7.2 to 6.8) or are shown descriptively without statistics (CD1; n = 3).

In a smaller cohort of *MTOR*^p.L2427P^ mice on the Swiss/CD1 background used by Ribierre *et al.*^19^ (n=3), continuous video-EEG monitoring was carried out under the same protocol. Seizures persisted throughout DQ treatment in this cohort as well (Figure 3A–D lower), consistent with the absence of an antiseizure effect in the C57BL/6J cohort. These data are shown descriptively, without statistics, because of low animal counts (average recorded EEG hours per CD1 animal: 900±100).

### Oral DQ Produced Only a Non-Significant Trend Toward Lower Senescence-Associated Markers Without Neuronal Loss

Although DQ neither eliminated cells nor altered seizures, the histological markers on which the senotherapy rationale rests could still have been affected, so we examined markers of senescence and mTOR activity in the FCD lesions (Figure 4A,B; DQ mice n=12, vehicle-only mice n=5). DQ densities are shown unscaled. Vehicle-only densities were multiplied by a single group-wide factor (1.47) equating their median tdTomato^+^ cell density to that of the DQ group, which DQ did not significantly alter (Figure 2F), to correct for differences in lesion size and IUE extent. Unscaled values are in *Source Data*. EGFP^+^ neuron density was unchanged (DQ: 94 [77–123] cells/mm²; Vehicle: 125 [107–141] cells/mm²; Mann–Whitney *U*=15, *P*=0.13, Bonferroni-corrected *P*=0.52; Figure 4C).

**Figure 4.**
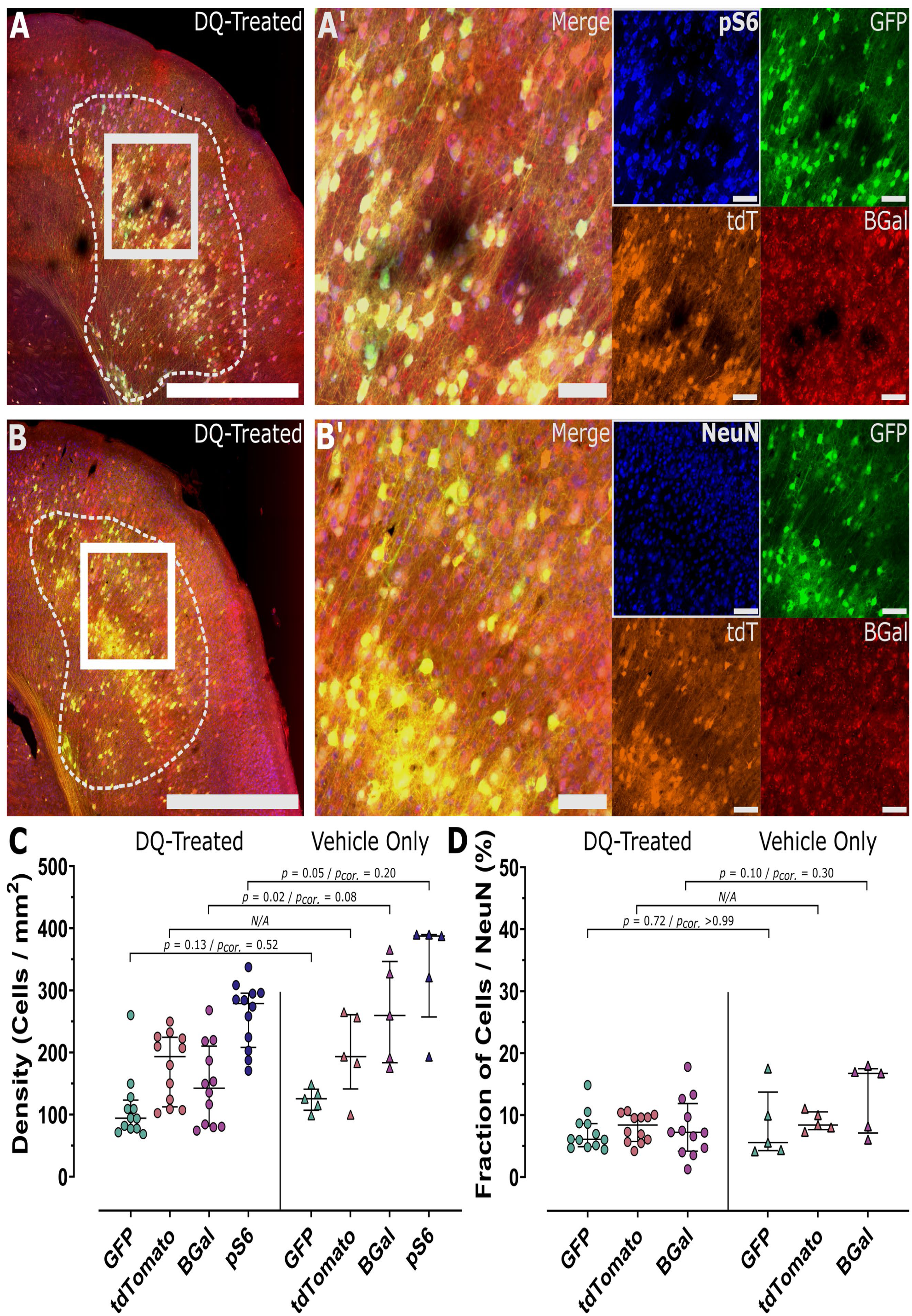
Histological markers of mTOR activity and senescence in FCD lesions after DQ or vehicle treatment. **A**, **B**) Representative confocal MIPs of coronal sections from the same DQ-treated mouse, with the FCD lesion outlined in white (dashed); boxes mark the regions enlarged in **A’** and **B’**. **A’**) Merged image and single channels for pS6 (blue), EGFP (green), tdTomato (orange; tdT) and B-Gal (red). **B’**) Merged image and single channels for NeuN (blue), EGFP (green), tdTomato (orange) and B-Gal (red). Scale bars = 1 mm (**A**, **B**) and 100 µm (**A’**, **B’**, including single channels). **C**) Spatial cell densities in FCD lesions of DQ-treated (circles; n = 12) and vehicle-only (triangles; n = 5) mice; each point is the mean of two brain sections from one animal; lines and error bars denote the median and IQR; vehicle-only densities were normalized to the median tdTomato^+^ density of the DQ group to correct for differences in lesion size and IUE lesion extent; two-sided Mann–Whitney *U* tests: *P*, uncorrected; *P*cor., Bonferroni-corrected across the four markers; median differences, DQ minus vehicle (Hodges–Lehmann, exact 95.2% CI, unadjusted, after vehicle scaling; cells/mm²): B-Gal −110 (−211 to −39); pS6 −95 (−150 to −13); EGFP −26 (−54 to 10); N/A, not tested. **D**) Cell densities as a fraction of NeuN^+^ cell density; display and group sizes as in **C**; *P*cor., corrected across the three markers.

**Figure 5.**
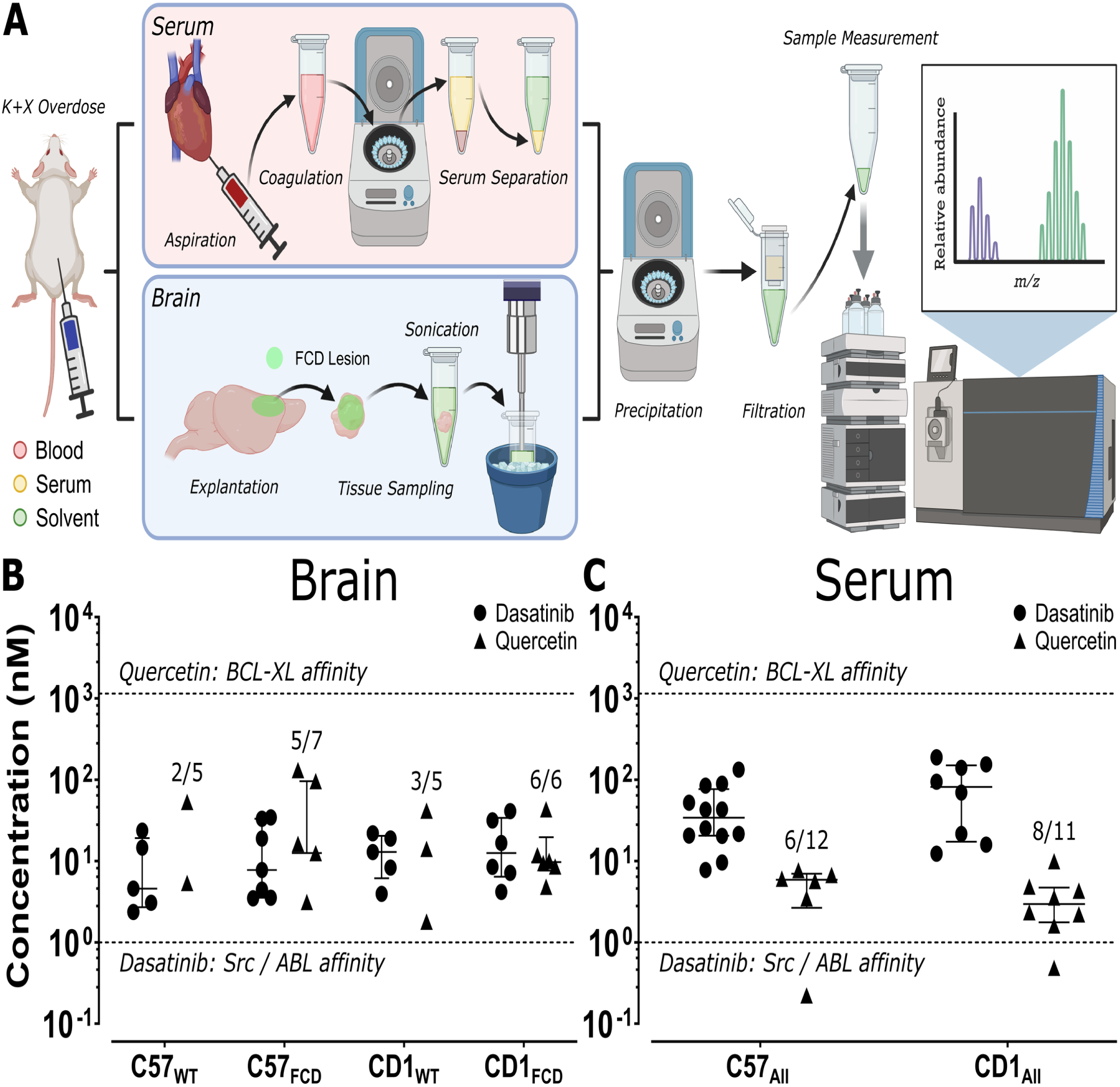
Dasatinib and quercetin concentrations in brain and serum measured by LC–MS/MS. **A**) Sample collection and processing. After ketamine/xylazine (K+X) overdose, blood was aspirated from the left ventricle and serum separated by coagulation and centrifugation; brains were then perfused with heparinized PBS, explanted, and homogenized by sonication; in FCD animals, brain tissue was sampled from the lesioned area as defined by EGFP fluorescence; in WT animals, brain tissue was sampled as the left cortical hemisphere. Extracts were protein-precipitated, filtered, and analyzed by LC–MS/MS with deuterated internal standards; tube colors denote blood (pink), serum (yellow), and extraction solvent (green). **B**) Brain and **C**) serum concentrations (nM, log scale) of dasatinib (circles) and quercetin (triangles); each point is one animal; lines and error bars denote the median and IQR where applicable. In **B**, groups are C57WT (n = 5), C57FCD (n = 7), CD1WT (n = 5), and CD1FCD (n = 6); dasatinib was quantified in every animal; fractions give the animals with quercetin above the lower limit of quantitation (LLOQ, 0.1 nM). In **C**, WT and FCD animals are pooled within strain (C57, n = 12 animals; CD1, n = 11); fractions give the animals with a valid quercetin measurement, and no valid serum value was below the LLOQ. Serum samples with absent or unreliable internal-standard signals were excluded (dasatinib, n = 3; quercetin, n = 9), leaving dasatinib n = 12 (C57) and n = 8 (CD1) and quercetin n = 6 (C57) and n = 8 (CD1). Dashed lines: reported *K*d of quercetin for BCL-XL (1.1 µM)^32^ (upper) and the upper bound of reported dasatinib IC50 values for Src-family kinases and ABL (1 nM)^31^ (lower). Subplot **A** was created with https://BioRender.com.

While B-Gal labeling showed a reduction in senescent cell density in the FCD lesion following DQ, this effect did not survive correction for multiple comparisons (DQ: 141 [81– 210] cells/mm²; Vehicle: 260 [183–346] cells/mm²; Mann–Whitney *U*=8, *P*=0.02, Bonferroni-corrected *P*=0.08; Figure 4C). Phosphorylated S6 (pS6), a downstream readout of mTOR activity, likewise showed no significant difference in spatial density between groups (DQ: 279 [208–296] cells/mm²; Vehicle: 388 [257–390] cells/mm²; Mann–Whitney *U*=11, *P*=0.05, Bonferroni-corrected *P*=0.20; Figure 4C).

Normalization to NeuN^+^ cell density (Figure 4B) did not reveal any latent B-Gal treatment effect attributable to differences in lesion size or neuronal packing density (Kruskal–Wallis *H*=6.244, *P*=0.28; Figure 4D). The fraction of B-Gal^+^/NeuN^+^ cells (DQ: 7.2 [4.2–11.9]%; Vehicle: 16.8 [7.1–17.5]%; Mann–Whitney *U*=14, *P*=0.10, Bonferroni-corrected *P*=0.30) indicates selective staining, with >80% of NeuN^+^ cells B-Gal-negative. These results indicate that oral DQ does not act as a senolytic that eliminates DNs.

### Brain Dasatinib Reaches Src/ABL-Inhibitory Concentrations, Whereas Quercetin Remains Far Below Its Reported BCL-XL Affinity

The non-significant trend toward lower senescence markers without cell loss raised the question of whether DQ reached the brain at concentrations consistent with its proposed mechanisms. We quantified dasatinib and quercetin by targeted LC–MS/MS with isotopically labeled internal standards (workflow in Figure 5A) in brain and serum from 23 mice sampled 90 min after the fourth daily dose, within the reported time-to-peak window for both compounds^28,29^ and practical for staggering animals. Groups were naïve C57BL/6J (n=5) and CD1 (n=5), C57BL/6J FCD (n=7), and CD1 FCD mice (n=6), completing a full strain-by-disease-status comparison, given reports of altered blood–brain barrier permeability in pathological cortex^30^. Dasatinib was quantifiable in every brain (median 8.5 nM, range 2.4–41 nM; Figure 5B) and in serum (median 34 nM in C57BL/6J and 82 nM in CD1; Figure 5C), with a brain-to-serum ratio of 0.22 (IQR 0.16–0.31; n=20 paired). Brain dasatinib did not differ by genotype or strain (stratified rank tests, *P*=0.38 for both), nor did serum dasatinib (genotype *P*=0.59; strain *P*=0.10). Brain dasatinib concentrations exceeded the Src/ABL IC_50_ (≤1 nM)^31^ at least two-fold.

Quercetin, in contrast, was measurable only at low nanomolar concentrations. It exceeded the limit of quantitation in 16 of 23 brains (C57_WT_ 2/5, C57_FCD_ 5/7, CD1_WT_ 3/5, CD1_FCD_ 6/6; Figure 5B), with a median of 12 [7.8–42] nM (range 1.8–131 nM) among quantifiable samples, far below its reported *K*_d_ for BCL-XL^32^ (Figure 5B). Because the median is computed only over samples above the limit of quantitation, it overestimates central tendency across the cohort; including the seven non-quantifiable brains would lower it further. Quercetin was quantifiable in serum in 14 of 23 animals (median 3.9 [2.2–6.0] nM, range 0.22–10 nM; Figure 5C); nine serum extracts were excluded because the internal-standard signal was absent or severely reduced. Among the 13 animals with quantifiable brain and serum values, brain quercetin exceeded serum quercetin in 11 (ratio 3.7, IQR 1.4–7.3), indicating that quercetin aglycone is not excluded from the brain and is present at low concentrations in both compartments. Quercetin was quantifiable in brain more often in FCD than WT mice (11/13 vs. 5/10; stratified by strain, exact permutation *P*=0.09), a difference we regard as exploratory.

## Discussion

Our multimodal dataset provides three main findings. First, longitudinal *in vivo* two-photon imaging directly demonstrates progressive loss of mTOR-hyperactive neurons in the FCD cortex (Figure 2C,D), extending prior cross-sectional histology to within-animal tracking. Second, lesions in this second, independently generated FCD II model contained B-Gal^+^ cells, a single marker consistent with, but not sufficient to establish, senescence (Figure 4A–C). Third, across all four modalities, we found no or limited evidence that DQ eliminates these cells or improves seizure outcome (Figures 2F, 3A–D, and 4C), in contrast to Ribierre *et al.*^19^; seizures likewise persisted in a smaller CD1 cohort of the same genotype (Figure 3A–D). One possible contributor is insufficient quercetin exposure, although this does not explain the difference from Ribierre *et al.*^19^. Dasatinib reached brain concentrations sufficient for substantial Src/ABL inhibition (Figure 5B), irrespective of FCD status, on a regimen matching the one originally reported^19^. Dasatinib exposure therefore appears adequate, although target engagement was not measured. Quercetin, by contrast, was quantifiable in only 16 of 23 brains, at a median of 12 nM (Figure 5B), nearly two orders of magnitude below the *in vitro K*_d_ of quercetin for BCL-XL (1.1 µM)^32^. Because quercetin, not dasatinib, is credited with the BCL-XL family-dependent apoptotic trigger, its low brain exposure offers a specific, testable hypothesis for the lack of senolysis. Several points bear on how this null should be read. It is not a question of sensitivity for the imaging endpoint. With nine animals, the paired design could detect about 16 additional percentage points of tdTomato⁺ loss during DQ (SD of differences 14.7; 80% power). The observed difference was 1.9 points (95% CI −9.4 to 13.2), which excludes the 70–80% elimination implied by the reported senolytic effect. For seizures, none of the seven C57BL/6J mice was seizure-free in the final recording window (exact 95% CI 0–41%), whereas seizure freedom was the reported outcome; the frequency comparison was less sensitive, detecting only changes of about 11 seizures/day (vehicle-phase mean, 3.3). Finally, the claims do not rest on equal evidence; elimination of mTOR-mutant neurons and the reported seizure freedom are excluded, whereas a senomorphic marker effect is not.

This interpretation is consistent with human pharmacokinetic data. In the first-in-human trial of DQ in early Alzheimer’s disease, dasatinib was detected in the cerebrospinal fluid (CSF) of four of five participants (CSF:plasma ratios 0.42–0.92%), whereas quercetin was undetected in CSF in any participant at any time point^20^. That the same asymmetry recurs across two mouse strains and in human CSF, the closest accessible proxy for brain exposure, suggests that low CNS exposure may be a property of oral quercetin rather than an artifact of our model. Quercetin was not, however, excluded from brain; it exceeded serum concentrations in 11 of 13 paired animals, so the limiting factor appears to be low systemic availability of free quercetin. Orally administered quercetin is almost entirely converted to glucuronide and sulfate conjugates during first-pass metabolism^33,34^, with free aglycone (the form required to engage BCL-2/BCL-XL) undetectable in human plasma and only trace in rodent plasma^33,34^. Our assay targets this free aglycone. In neuronal-like SH-SY5Y cells, quercetin at 2.5–5 µM attenuated stress-induced senescence markers without reducing cell number^35^, far above our median brain level.

Soma enlargement is a core mTORC1-dependent feature of DNs. Rapamycin reduced soma size and seizures in the same *MTOR*^p.L2427P^ model when started after seizure onset^21^ and prevented dysmorphogenesis in a Rheb-driven model when given from birth, with soma size returning after withdrawal^36^. DQ, in contrast, did not change soma diameter, and pS6^+^ density did not differ significantly after correction, consistent with DQ not acting through mTORC1 suppression.

Our null result is not isolated, and DQ’s senolytic efficacy has repeatedly failed to generalize. DQ failed to clear senescent cells and appeared acutely pro-tumorigenic in hepatocellular carcinoma^37^, increased rather than reduced kidney damage biomarkers in acute kidney injury^38^, and failed to improve the antiviral immune response in aged influenza-challenged mice^39^. Most directly relevant, a recent report found that DQ, at doses at or below those used here, induced demyelination in the corpus callosum of healthy young and aged mice without oligodendrocyte death, with oligodendrocyte stress responses proposed as a mechanism^40^. This is a functional effect on a CNS cell population without measurable cell loss.

This study is not a direct replication. We used a different *MTOR* variant, a different animal background for the longitudinal cohort, a different reporter system, and denser seizure ascertainment. Whether a senolytic effect generalizes beyond the model in which it was characterized bears directly on its clinical relevance, and no single agent is expected to work across all FCD II mutations. One explanation for the divergence is that the two variants are not interchangeable. S2215F and L2427P are both recurrent gain-of-function substitutions in patients^19,21,41^, but S2215 lies at the interface between the N- and C-lobes of the kinase domain, whereas L2427 lies in the kα9b-helix, which together with the negative regulatory domain occludes the catalytic cleft^42^. Modeling of the oncogenic L2427R substitution predicts cleft opening, and the same authors expect S2215 to act similarly^42^, so any difference would have to lie in downstream signaling, which has not been compared directly. Such a difference is plausible, since whether a cell depends on MCL-1 or on BCL-2/BCL-XL for survival varies with context, including mTORC1 status^43^. A recent preprint reports a profile “fundamentally distinct” from TSC2 loss for an overgrowth- syndrome gain-of-function *MTOR* mutation^44^. This evidence compares pathway nodes and cell types, not distinct alleles of *MTOR*; whether allele-level divergence exists is untested, although different *MTOR* mutants do differ in mTORC1 potency and in disruption of cortical migration^45^.

Independently of exposure, and even if CNS concentrations were adequate, a second question is whether the senescent DNs in our model are susceptible to quercetin at all. If L2427P engages this landscape differently from S2215F, senescent DNs in our model could depend on a different balance of anti-apoptotic proteins. Quercetin’s senolytic mechanism is itself not uniform. It binds the BH3 domain of BCL-2 and BCL-XL (*K*_d_ 1.1 µM for BCL-XL^32^), but has also been reported to lower MCL-1 in leukemia cells at tens of micromolar levels^46,47^. This supports a testable hypothesis for follow-up: profiling the anti-apoptotic dependency of L2427P- and S2215F-driven senescent neurons directly, for example by BH3 profiling, which predicts senolytic response more reliably than genotype or marker expression alone^48^. Our histology (Figure 4C) shows senescence markers with only a non-significant trend toward lower B-Gal density after DQ, consistent in direction with a senomorphic effect. As no marker comparison survived correction, this remains hypothesis-generating and does not identify the underlying dependency.

Beyond DQ itself, several paths follow. One is to reformulate quercetin or to substitute a BCL-2 family-targeting agent with better native CNS penetration. Another is to accept a senomorphic strategy, for which brain-penetrant mTOR inhibitors are in early development^49^; these have improved CNS penetration relative to the rapalogs already used clinically^8,9^. Clinically, these results argue against assuming that a senolytic effect shown in one FCD II genotype will hold in another and for demonstrating brain exposure of each component before translation.

### Limitations

This study has several limitations. Treatment order was not counterbalanced, as vehicle always preceded DQ against a background of progressive neuronal loss. The vehicle-only group controls for this at the group level, and because loss accumulates with time, any residual bias would act against, rather than in favor of, a null DQ effect. Non-FCD controls differ in genotype and labeling method and were imaged on a less dense schedule (four sessions over six weeks versus six over eight), so they bound gross imaging drift rather than serving as a matched control. The internal control for the loss measurement is that the tdTomato^+^ and EGFP^+^ populations declined together in the same animals. Our CD1 cohort is limited to EEG and LC–MS/MS endpoints, because no Cre-dependent tdTomato reporter line exists on the outbred CD1 background and AAV labeling proved too inconsistent for confident quantification. BCs, which also carry the senescence signature in human FCD IIb tissue^19^, are not modeled by IUE. We did not assess white matter integrity in DQ-treated tissue^40^, nor characterize the anti-apoptotic dependency of senescent DNs directly. Our sample sizes for EEG (n_C57_=7; n_CD1_=3) and LC–MS/MS (n=5– 7 per group) are modest, and no histological comparison survived correction for multiple comparisons. Senescence was assessed with a single marker (B-Gal) and senolytic readouts were obtained only in C57BL/6J L2427P mice; S2215F was not tested. DQ began at about week 20, histology followed about two weeks after the last dose, and the post-treatment EEG window was two weeks, versus 16 weeks of age, one month, and four weeks, respectively, in Ribierre *et al.*^19^. Drug levels were total, not unbound, measured once after four doses with an unvalidated assay, and target engagement (for example pSrc or pCrkL) and a positive-control senolytic were not assessed. Cre recombination is incomplete, so some mutant neurons may be tdTomato-negative and contribute to the EGFP-only population. A few thousand mutant neurons suffice to trigger seizures in this model^22^.

## Conclusion

This study makes two contributions. Methodologically, it establishes, to our knowledge, the first longitudinal *in vivo* imaging platform for tracking individual mTOR-mutant neurons in a mouse model of FCD, directly demonstrating progressive neuronal loss that cross-sectional histology could only infer. Pharmacologically, we used that platform to ask whether senescent-cell clearance by oral DQ generalizes beyond the model in which it was characterized. Cellular senescence remains a compelling candidate driver of epileptogenesis in FCD II and our data corroborate its presence across independent models. The reported senolytic effect, however, was not reproduced in our model, and our pharmacokinetic data point to one possible reason: quercetin’s apparent failure to reach concentrations in brain at which its senescence-modulating effects are seen *in vitro*, further supported by a human cohort. A second possibility is that the *MTOR* variant shapes which anti-apoptotic pathway senescent DNs depend on. Neither is evidence against senotherapy in FCD as a concept, and both are evidence that the pharmacological warhead and the driving mutation are important experimental variables.

## Acknowledgments

The authors would like to thank Pavlína Smolková for plasmid preparations (amplification).

## Funding

This work was supported by the Ministry of Health of the Czech Republic (grants NW24-04-00041 and NW26-04-00517); the Ministry of Education, Youth and Sports of the Czech Republic (project LX22NPO5107, National Institute for Neurological Research, funded by the European Union – Next Generation EU; and project Brain dynamics, OP JAK, CZ.02.01.01/00/22_008/0004643, co-funded by the European Union); and Charles University (project EXCITE, UNCE24/MED/021; PRIMUS/23/MED/011; and GA UK grant No. 423325). The funders had no role in the design of the study, the collection, analysis, or interpretation of data, the writing of the manuscript, or the decision to submit it for publication.

## Author Contributions

According to the CRediT taxonomy, the contributions of individual authors were as follows: Conceptualization: CVLO, BS, and ON. Methodology: CVLO, NS, and JK. Software: CVLO, JTR, and JK. Formal analysis: CVLO, NS, KČ, JP, SW, BR, BK, and JK. Investigation: CVLO, NS, NP, KČ, MŘ, JP, SW, BR, BK, and ON. Data curation: CVLO, NS, NP, and KČ. Writing: CVLO, BS, PJ, and ON. Visualization: CVLO. Supervision: PJ and ON. Project administration: CVLO, PJ, and ON. Funding acquisition: JP, JK, PJ, and ON.

## Conflict of Interest Statement

None of the authors has any conflict of interest to disclose. We confirm that we have read the Journal’s position on issues involved in ethical publication and affirm that this report is consistent with those guidelines.

## Data Availability Statement

The data that support the findings of this study are available in the supplementary material of this article under *Source Data*. Raw data files, including raw imaging data (tens of terabytes), are available from the corresponding author upon reasonable request.

## Ethics Statement

All animal care and use conformed to institutional policies and guidelines. Experiments were performed in accordance with the Animal Care and Animal Protection Law of the Czech Republic, which is compatible with European Union Directive 2010/63/EU, and the protocols were approved by the Ethics Committees of the Second Faculty of Medicine, Charles University (Project License No. MSMT-26023/2023-5). The principles outlined in the ARRIVE guidelines and the Basel Declaration, including the 3R concept, were considered when planning the experiments. Anesthesia, analgesia, and terminal sampling are described in *Extended Methods*.

### Replacement

the fate of individual neurons, seizures, and brain drug exposure in an epileptogenic cortex could not be answered *in vitro*.

### Reduction

in the longitudinal C57BL/6J cohort, each animal served as its own control and contributed imaging, video-EEG, and histology data, so that 17 animals produced all longitudinal datasets; pharmacokinetic sampling used separate terminal cohorts in which each animal was used once, and the four non-FCD imaging controls were animals primarily used in a different project. The CD1 EEG cohort was kept small and analyzed descriptively. The number of animals contributing to each analysis differs because of attrition and the requirement for complete paired data across sessions (Table 1); no animal was excluded on the basis of its result, individual pharmacokinetic samples failing internal-standard criteria were excluded for that analyte only, and 47 animals were used in total.

### Refinement

surgeries were performed under isoflurane anesthesia, with ketoprofen (5 mg/kg, subcutaneously, before surgery and daily for two days afterward) given for *in utero* electroporation and for cranial window and electrode implantation; animals were group-housed under a 12/12 h light/dark cycle with food and water *ad libitum*; terminal sampling followed a ketamine/xylazine overdose (120 and 20 mg/kg).

## Notes

### Competing Interest Statement

The authors have declared no competing interest.

### Summary of Updates

Version 2 is a substantially expanded, full-length version of the study. The LC-MS/MS pharmacokinetics have been considerably extended, from 8 to 23 animals in a complete strain-by-disease-status design (C57BL/6J and Swiss/CD1, wild-type and FCD), with both dasatinib and quercetin now quantified in brain and serum and compared by stratified rank tests, and continuous video-EEG seizure monitoring has been added in a Swiss/CD1 cohort carrying the same MTOR variant. The manuscript has been rewritten and expanded throughout, with new figures, full Introduction, Methods, Discussion and Limitations sections, and additional statistics including effect sizes with confidence intervals and post hoc sensitivity analyses.

