## Supplementary Methods for "Chronic *In Vivo* Imaging of a Focal Cortical Dysplasia Mouse Model Reveals Progressive Neuronal Loss with No Senolytic or Antiseizure Efficacy Following Dasatinib–Quercetin Treatment"

### Extended Methods

#### Sample Preparations

##### *Animals*

Animals of either sex were housed in groups under standard conditions in a room with controlled temperature (22±1 °C) and a 12/12 h light/dark cycle and were given chow and water *ad libitum*. Experimental C57BL/6J FCD mice (n=24) were generated by breeding wild-type (WT) C57BL/6J (strain #000664; Jackson Laboratory, Bar Harbor, USA) with the Cre-dependent LSL-tdTomato Ai9 reporter line (strain #007909; Jackson Laboratory): 17 entered the longitudinal protocol (12 DQ-treated, 5 vehicle-only) and contributed to the imaging, EEG, and immunohistochemistry datasets (the five vehicle-only animals contributed histology only), and a further 7 formed a separate terminal cohort for LC–MS/MS. CD1 FCD mice (n=9) comprised 3 for continuous video-EEG and 6 for a separate terminal LC–MS/MS cohort. WT C57BL/6J mice (n=5) and WT CD-1/ICR mice (strain #022; Charles River, Wilmington, USA; n=5) were used for LC–MS/MS only. Four mice on a mixed C57BL/129S4 background (>75% C57BL/6J), generated by crossing VIP-IRES-Cre (strain #010908; Jackson Laboratory) to Ai9, served as non-FCD imaging controls with neurons labeled by AAV-delivered GCaMP7f. In total, 47 animals were used. Because attrition and the requirement for complete paired data across sessions meant that not every animal contributed to every analysis, the *n* per analysis is given in *Table 1*; no animal was excluded on the basis of its result. Individual pharmacokinetic samples that failed internal-standard criteria were excluded for that analyte only. Both males and females were included. Animals were randomly allocated to the DQ or vehicle-only groups. Within animals, the vehicle phase always preceded DQ and was therefore not randomized. No formal sample-size calculation was performed, and group sizes reflected prior work and animal-use constraints (3R).

All animals were housed in the same room, and experiments were staggered over many weeks according to the weekly yield of electroporated litters. The person aware of group allocation did not perform the two-photon imaging or the EEG seizure labeling. Body weight ranged from 25 to 30 g in C57BL/6J mice and from 35 to 40 g in CD1 mice. Criteria for inclusion in the imaging and EEG analyses were set before the study: a clear cranial window, adequate EEG signal quality, and a seizure phenotype. Failure to meet these criteria was the reason for attrition from the 17 animals in the longitudinal cohort to those analyzed for imaging and EEG. Animals were assessed daily and were imaged or recorded only if bright, alert, and responsive. Mice had a hiding house, and before the first imaging session each mouse had one 20-min session of acclimatization to head fixation. No adverse events occurred. At the end of the experiments, all animals were killed by ketamine/xylazine overdose.

All experiments were performed under the Animal Care and Animal Protection Law of the Czech Republic, fully compatible with the guidelines of the European Union directive 2010/63/EU. The protocol was approved by the Ethics Committees of the Second Faculty of Medicine (Project License No. MSMT-26023/2023-5).

##### *In Utero Electroporation and FCD Evaluation*

The FCD lesion was induced by *in utero* electroporation (IUE). Pregnant mice (day 14.5±0.5 post-fertilization) were anesthetized with isoflurane (4% induction, 1.5–2%

maintenance, 0.5 L/min O2), subcutaneously injected with ketoprofen (5 mg/kg), and positioned on their backs. A midline incision was performed along the *linea alba*. The uterine horns were gently exposed. Using pulled and sharply beveled glass capillaries, the left lateral ventricle of each embryo was injected with a mixture of plasmids dissolved in ACSF containing 2.0 µg/mL Fast Green (F7252; Sigma-Aldrich, St. Louis, USA). For FCD model animals (n=33), the plasmid mixture contained 3.0 µg/µL mutant *mTOR* (p.L2427P; SoVarGen, Daejeon, Korea) followed by tamoxifen-activated CreERT2, all under the control of the CAG promoter. For the lesion visualization, the mTOR plasmid was mixed with an expression plasmid CAG-EGFP at a final concentration of 1.5 µg/µL.

After each injection, the electroporation was performed by positioning forceps-type electrodes (3 mm platinum plated pads, CUY650P3; Nepa Gene, Chiba, Japan) on the head of the embryo, and applying five 35–45 V, 50 ms pulses with 950 ms interpulse intervals using a custom-made high-resolution electroporator with a 40 mA current limiter[1]. The uterine horns were returned to the abdominal cavity and the wound was sutured in individual layers. The mouse was turned on its abdomen and kept on the inhalation mask with pure oxygen until regaining consciousness. Then, the mouse was returned to its home cage that was placed on a heating pad overnight. Following surgery, the mice were injected daily with ketoprofen (5 mg/kg) for two consecutive days. When the pups were born, we screened them for fluorescence using a strong, collimated blue LED (GTP04B3410320; Shenzhen Getian Opto-electronics, Shenzhen, China) with a bandpass (BP) excitation filter (475/50 nm; Edmund Optics, Barrington, USA) and a stereomicroscope (Model STM 823 N; Optika, Ponteranica, Italy) with a large-diameter lens cap containing a green filter (BP 525/50 nm; Edmund Optics). Only the successfully electroporated pups in which we detected a properly positioned green fluorescent area were kept in the experiment. Litters with heterogeneous or bilateral fluorescent lesion patterns were excluded.

CD1 FCD mice were generated by IUE of WT Swiss/CD1 embryos with the same CAG-mTOR$^{p.L2427P}$-IRES-CreERT2 and CAG-EGFP plasmid mixture used for the C57BL/6J cohort; the *MTOR* construct was kept identical for consistency and EGFP was co-electroporated both to screen pups for successful electroporation and to delineate lesion extent. Because no Cre-dependent LSL-tdTomato reporter line is available on the outbred Swiss/CD1 background, mTOR-mutant neurons could not be genetically labeled as in the C57BL/6J cohort; AAV-mediated labeling was attempted instead, but signal appearance and stability proved too variable for confident longitudinal quantification, so two-photon imaging and immunohistochemistry are not reported for this cohort. CD1 FCD mice were therefore used for continuous video-EEG (n=3) and, in a separate terminal cohort, for LC–MS/MS blood and brain sampling (n=6).

##### *Tamoxifen Administration*

C57/LSL-tdTomato mice which received mTOR-CreERT2 were later given tamoxifen via oral gavage at W5 of age. Tamoxifen (#T5648; Sigma-Aldrich) was dissolved in corn oil (#C8267; Sigma-Aldrich) at a concentration of 20 mg/mL by stirring at 60 °C for 20 min and protected from light. Mice received tamoxifen by oral gavage at a dose of 3 mg per animal (roughly 100 mg/kg for a 30 g mouse) once daily for five consecutive days[2].

##### *Cranial Window and Electrode Implantation Surgery*

Mice were operated on at 12–13 weeks of age. Animals were anesthetized with isoflurane in oxygen (4% induction, 1.5–2% maintenance, 0.5 L/min O2), subcutaneously injected with ketoprofen (5 mg/kg) before surgery and daily for the following two days (three doses in total), and placed on a 37 °C heated pad. The fur on top of the animal's head was shaved, and a small area of skin was removed to expose the dorsal skull. A ring-shaped titanium headbar with side wings and a preassembled video/EEG connector was glued over the left parietal cortex using a gel form of cyanoacrylate (CA) glue (Loctite #1363589; Henkel, Düsseldorf, Germany) and cured with a glue accelerator (Insta-Set; Bob Smith Industries, Atascadero, USA). We carefully performed a 4 mm craniotomy over the FCD area using a handheld high-speed microdrill (Success40; Osada Electric, Tokyo, Japan) and then gently removed the bone flap to expose the intact dura. The craniotomy was then closed with a 4 mm round #1.5 cover glass that was fixed to the skull with CA glue. We implanted two LFP electrodes: one placed directly over the FCD by the edge of the cover glass, the second one to the contralateral cortex over the homotopic area. The grounding/reference electrodes were placed over the inferior colliculus/cerebellum. The skin margins were secured and sealed to the headbar/connector construct and the skull using CA glue and the area was disinfected.

##### *AAV Injection Surgery*

Four non-FCD animals, primarily used in a different project, were used as controls for *in vivo* imaging and underwent a similar cranial window surgery, including the same analgesia. In this procedure, we additionally injected several sites of the cortex with small volumes of an adeno-associated virus (2×30 nL of $1\times10^{11}$ GC/mL AAV9-hSyn-jGCaMP7f) to a depth of 250 μm below the dura. For atraumatic injection, we used pulled and beveled glass capillaries (30° angle, 10–15 μm outer diameter; 3-000-203-G/X; Drummond Scientific, Broomall, USA) and injected at a rate of 1 nL/s. A Nanoject III nanoinjector (Drummond Scientific) was used to drive the glass micropipette that was held in place for 5 min to prevent the virus from backleakage. The craniotomy was then closed with a 4 mm round #1.5 cover glass that was fixed to the skull with CA glue.

##### *Senolytic Administration Protocol*

Dasatinib (#HY-10181; MedChemExpress, Monmouth Junction, USA) and quercetin (#Q4951; Sigma-Aldrich) stock solutions were prepared in DMSO (#472301; Sigma-Aldrich) one day before protocol initiation at concentrations of 48 mg/mL and 200 mg/mL, respectively, aliquoted, and stored at −20 °C. The final gavage solution was prepared immediately before administration to achieve concentrations of 12 mg/kg dasatinib and 50 mg/kg quercetin in 30% PEG300 (#91462; Sigma-Aldrich) and 5% Tween-20 (#P1379; Sigma-Aldrich) in ddH2O, aiming for an administered volume of 0.15 mL. Mice received vehicle (10% DMSO in PEG300/Tween-20/ddH2O) or senolytic treatment by oral gavage for four consecutive days, followed by a two-day intermission, and an additional five consecutive days of administration, per the specific phase of the experimental protocol.

##### *Brain Tissue Slice Preparation*

After the completion of the longitudinal experiments, mice were overdosed with an intramuscular injection of a mixture of ketamine/xylazine 120 mg/kg and 20 mg/kg, respectively. The animals were transcardially perfused with cold phosphate-buffered saline (PBS) and 4% paraformaldehyde (PFA). Brains were collected, post-fixed in PFA overnight, and then stored in PBS with 0.05% azide. To further map the position and quantify the spatial extent of the lesion, the post-fixed brains were photographed using the aforementioned blue excitation/green emission add-on to the surgery microscope. The fixed brains were cut on a vibratome (Leica VT1200 S; Leica, Wetzlar, Germany) into 60 µm thick coronal slices and stored in PBS with 0.05% azide.

##### *SaβGal Colorimetric Assay*

Once all brains had been sliced, slices were washed twice with PBS + 1 mM $MgCl_2$ buffered to pH 6.0 with HCl (PBS/$MgCl_2$). Following the second wash, an X-Gal staining solution containing 2.5% 0.12 mM $K_3Fe(CN)_6$, 2.5% 0.12 mM $K_4Fe(CN)_6 \cdot 3(H2O)$, and 2.5% X-Gal (#15520018; Thermo Fisher Scientific, Waltham, USA) in PBS/$MgCl_2$ was applied to the free-floating slices. The slices were then incubated with gentle shaking at 40 °C for 14 h. All slices were processed together in the exact same experimental conditions, such as pH, temperature, humidity, and incubation time. Afterwards, the slices were rinsed twice with fresh PBS and stored for subsequent processing. We prepared at least six slices from each FCD mouse.

##### *Immunohistochemistry*

The free-floating slices were washed with PBS and non-specific binding sites were blocked using ChemiBLOCKER (#2170; MilliporeSigma, Burlington, USA). Immunohistochemistry (IHC) was performed using primary antibodies for neuronal nuclear protein (NeuN; 1:1000, rabbit; #ABN78; Sigma-Aldrich) and phospho-S6 ribosomal protein (Ser235/236) (1:600, rabbit; #2211S; Cell Signaling Technology, Danvers, USA). All primary antibodies were incubated with the slices overnight at 4 °C. We used a secondary antibody conjugated with a fluorescent dye CF 405S (1:500, anti-rabbit; #SAB4600024; Sigma-Aldrich). Secondary antibodies were also incubated overnight at 4 °C. After washing, slices were mounted using Fluoromount-G Mounting Medium (#00-4958-02; Thermo Fisher Scientific). We prepared at least three slices from each FCD mouse with each antibody.

##### *Tissue Lysates and Compound Extraction*

Low-molecular-weight (LMW) compounds were extracted from mouse serum and brain tissue using a cold organic precipitation method. Acetonitrile and methanol (LC–MS grade) were mixed 1:1 (v/v) and supplemented with 0.1% formic acid to generate the extraction solvent, which was pre-chilled to −20 °C before use. A second batch of extraction solvent was prepared to the same specification with the addition of two deuterium-labeled internal standards (IS) at a concentration of 200 nM: dasatinib-d8 (#HY-10181S; MedChemExpress) and quercetin-d3 (#HY-18085S1; MedChemExpress). All animal samples were extracted with the IS solution except two, which were used to create matrix calibration curves. Tissues were collected 90 min following the final gavage.

**Serum Samples.** Mice (n=23) received the first four daily doses of the gavage regimen and, 90 min after the fourth dose, were overdosed with an intramuscular injection of a mixture of ketamine/xylazine 120 mg/kg and 20 mg/kg, respectively. Whole blood (0.5–1.0 mL) was aspirated from the left ventricle using a 20GA needle and syringe and transferred to uncoated tubes. The blood was then allowed to coagulate at room temperature for 10–15 min at which point tubes were centrifuged at 2000 × g for 10 min at 4 °C. Serum was carefully transferred to fresh tubes without disturbing the pellet. 300 µL aliquots of serum were transferred to pre-chilled microcentrifuge tubes and three volumes of cold extraction solvent were added. Samples were vortexed briefly and incubated at −20 °C for 20 min to ensure complete protein precipitation. Samples were then centrifuged at 16,000 × g for 10 min at 4 °C. Clarified supernatant was transferred to 0.22 μm filter tubes (#CLS8160; Sigma-Aldrich) and centrifuged at 16,000 × g for 15 min at 4 °C. Finally, the filtered liquid was transferred to 30 kDa MWCO filter tubes (#MRCF0R030; MilliporeSigma) and centrifuged at 16,000 × g for 15 min at 4 °C. All filtered samples were then collected and stored at −80 °C until use.

**Brain Samples.** Following the blood aspiration step from above, mice (n=23) were perfused with 20 mL of ice-cold PBS with heparin (20 IU/mL). Freshly explanted brains were weighed (230±6 mg wet weight, mean±SEM) and immediately homogenized on ice using a sonicator (Sonoplus UW 2070; Bandelin Electronic, Berlin, Germany) in pre-chilled extraction solvent at a ratio of 3 μL solvent per 1 mg tissue. Homogenates were incubated at −20 °C for 20 min to facilitate protein precipitation. All steps following protein precipitation were the same as for serum samples above.

**Calibration Curves.** Two animals were processed using a non-IS extraction solvent mixture for the generation of calibration curve points. These animals had not received DQ. Serum and brain extracts were pooled from these animals to create the background matrix. Final calibration points were created from serum and brain samples at concentrations ranging from 0.1–1000 nM for dasatinib and quercetin, separately, and fixed concentrations of dasatinib-d8 and quercetin-d3 at 100 nM. Terminal sampling was performed 90 min after the final gavage, within the reported time-to-peak window for both compounds, so that measured concentrations approximate peak rather than trough exposure. Quercetin additionally shows a second absorption peak attributable to enterohepatic recirculation[3], so a single terminal sample cannot characterize total exposure across the dosing interval.

#### Data Acquisition

##### In Vivo *Two-Photon Imaging*

For intravital two-photon imaging, we used a custom-made high-resolution two-photon microscope. The microscope was equipped with a tunable femtosecond laser (Chameleon Discovery NX TPC; Coherent, Saxonburg, USA). Laser power and blanking were controlled within the laser head. The angular range of the scanner engine (Resonant Scan Box; Sutter Instrument, Novato, USA) was adjusted to produce a square field of view (FOV) of 810 μm at 1× zoom when using a 20× high-NA objective (XLUMPFLN 20×, NA 1.0, WD 2.0; Olympus, Tokyo, Japan).

Fluorescence emission was separated by deflection by primary dichroic (ZT670rdcxxrxt-UF1; Chroma Technology Corp., Bellows Falls, USA) and collected through a 2″ detection pathway. The emission was spectrally separated into green and red channels by a secondary dichroic mirror (T570lpxr; Chroma Technology Corp.), bandpass filtered (Green: ET525/50m; Red: ET605/52m; Chroma Technology Corp.), and detected by two cooled silicon photomultipliers (PDA43; Thorlabs, Newton, USA). The signals were digitized/recorded using an NI-5732 digitizer connected to an NI-PXIe-7961 FPGA and NI-PXIe-1073 chassis (National Instruments, Austin, USA). Data acquisition, scanning, and laser control were performed using the ScanImage software suite (version 2020.1.4; MBF Bioscience, Williston, USA).

Four weeks after surgery, mice were placed in a custom-made holder and head-fixed via the titanium headbar. A custom-made laser-reflection device was used to precisely level the cranial window surface to the horizontal plane. This step is essential for repeated imaging across longitudinal sessions to ensure relocation of the same neuronal population. Each mouse contributed one image sampling site.

To ensure reliable relocation of the same region of interest across longitudinal sessions, a set of reference images was collected during the first week of imaging: (*i*) an overview of the cortical surface with vascular landmarks and (*ii*) an image at the target imaging depth (Z-depth = 0 µm). These reference images were used in all subsequent sessions to consistently identify the same recording sites. Imaging was performed at Z-depths ranging from +60 µm to −190 µm relative to 0 µm. Laser power was kept at <50 mW under the objective using 1020 nm excitation. Image sequences were acquired at 60 frames per Z-slice with a Z-step size of 1.6 µm and an image resolution of 1024×1024 pixels.

##### *EEG Recording*

Following at least a one-week recovery period after surgery, mice (n=7) were individually video-EEG monitored continuously for ten weeks. Custom-made video-EEG units consisted of a counterbalance mechanism, slip rings with an active motorized commutator, and a video camera. The connector on the head of the animal was connected to a headstage in which the spontaneous electrographic activity was amplified, band-pass filtered (0.1 Hz–1.6 kHz), and sampled at 5 kHz using a 32-channel headstage amplifier with an AD converter (#RHD2132; Intan Technologies, Los Angeles, USA). The data were recorded using an interface board (#C3100; Intan Technologies) controlled by Spike2 software (version 10.02; Cambridge Electronic Design, Cambridge, UK). Manual seizure labeling and EEG data analysis were performed using custom-made scripts in a MATLAB 2019b computing environment (MathWorks; Natick, USA).

CD1 FCD mice (n=3) were monitored under the same protocol, yielding an average of 900±100 recorded EEG hours per animal (mean±SEM). Owing to the small cohort size, CD1 seizure data are reported descriptively, without statistical comparison.

An electrographic seizure was defined as a run of rhythmic epileptiform activity of amplitude at least 2–3× baseline, dominant frequency above 2 Hz, lasting at least 10 s, showing evolution in amplitude and frequency and returning to baseline at offset. Events separated by less than 10 s were counted as one. Recordings were scored manually and blind to recording phase. Behavioral correlates were verified from synchronized video and scored on a modified Racine scale[4,5]; seizures were classified as non-convulsive (stages 1–2) or convulsive (stages 3–5). Recording and scoring followed AES/ILAE Translational Task Force standards for rodent video-EEG[6].

##### *Confocal Microscopy*

Images from coronal brain slices were captured using a high-resolution spinning-disk confocal, Andor Dragonfly 503 (Oxford Instruments, Abingdon, UK). The Dragonfly 503 microscope was equipped with a HC PL APO 20×/0.75 NA IMM CORR CS2 multi-immersion objective (Leica), a quad excitation dichroic filter set (405/488/561/640 nm), and a Zyla 4.2 Plus sCMOS camera (Oxford Instruments). We imaged in four color channels with WLL illumination with the following Zyla emission filters: blue (BP 450/50 nm), green (BP 525/50 nm), red (BP 600/50 nm) and far-red (BP 700/75 nm). The final confocal voxel tiles were sampled at 0.301×0.301×1.480 µm voxels.

##### *Liquid Chromatography–Mass Spectrometry*

Brain tissue concentrations of dasatinib and quercetin were quantified by liquid chromatography–tandem mass spectrometry (LC–MS/MS). Chromatographic separation was performed on an ExionLC AD system with autosampler (SCIEX, Framingham, USA) using a Kinetex F5 column (1.7 µm, 100×2.1 mm, 100 Å; Phenomenex Inc., Torrance, USA) with a SecurityGuard ULTRA F5 pre-column cartridge (2.1 mm ID). The column was maintained at 40 °C with an autosampler temperature of 5 °C and an injection volume of 2 µL. Mobile phase A consisted of deionized water (Milli-Q Integral 3; MilliporeSigma) with 0.1% formic acid (LC–MS grade; Thermo Fisher), and mobile phase B consisted of methanol (HiPerSolv Chromanorm; VWR International, Leuven, Belgium) with 0.1% formic acid, delivered at 0.2 mL/min using the following gradient: 80% A / 20% B from 0–1.5 min, ramping to 85% B at 8.0 min, then to 100% B at 8.6 min, held at 100% B until 11.5 min, returning to 80% A / 20% B at 11.6 min and held isocratically until 15.0 min.

Mass spectrometric detection was performed on a Triple Quad 6500+ system (SCIEX) using electrospray ionization (ESI) in multiple reaction monitoring (MRM) mode. Dasatinib was detected in positive ESI mode (ion spray voltage +5.5 kV, source temperature 350 °C, curtain gas 40 psi, GS1/GS2 50 psi) monitoring the transition m/z 488.1 → 401.0 (collision energy 39 V) as the quantifier and m/z 488.1 → 232.0 (55 V) as the qualifier, with a declustering potential of 246 V and dwell time of 30 ms. Quercetin was detected in negative ESI mode (ion spray voltage −4.5 kV, entrance potential −10 V) monitoring the transition m/z 300.9 → 150.8 (collision energy −30 V) as the quantifier and m/z 300.9 → 178.9 (−24 V) as the qualifier, with a declustering potential of −85 V. Deuterium-labeled internal standards dasatinib-d8 and quercetin-d3 were used for quantitation of dasatinib (m/z 496.2 → 406.0, quantifier) and quercetin (m/z 303.9 → 150.7, quantifier), respectively.

Calibration curves were prepared in brain tissue matrix over a concentration range of 0.1–1000 nM and fitted using a linear regression model with $1/x^2$ weighting. Blank matrix calibrations for dasatinib spanned 0.1–1000 nM and were fitted with a quadratic regression model with $1/x^2$ weighting. The lower limit of quantitation (LLOQ) was defined as the lowest concentration of the respective calibration curve, i.e., 0.1 nM. Quality control samples were prepared from independently spiked brain homogenate and processed alongside study samples; no additional formal validation including inter/intra-day precision, matrix effect assessment, or recovery determination was performed. No dilution steps were applied prior to injection.

#### Data Processing

##### *Two-Photon Image Stacks*

Z-stacks from longitudinal sessions were aligned to the first (reference) imaging session using a rigid transformation using ITK-elastix in a Python environment[7,8]. Raw two-photon image stacks were acquired in ZCYX format (Z-planes × 2 channels × Y × X) and processed using a custom Python pipeline employing the tifffile and NumPy libraries. To correct for bidirectional scanning artifacts, phase correlation was applied between the mean intensity profiles of even and odd rows from the central Z-plane, averaged across channels, and the estimated pixel shift was used to apply a compensatory row-wise correction to all odd rows across the full stack, with edge-value padding at row boundaries. To account for Z-drift across imaging sessions, stacks were cropped about a manually determined Z center rather than the geometric midpoint. Z centers were identified for each timepoint by visually inspecting the raw stacks in FIJI (ImageJ 2.16.0/1.54p)[9] and scrolling through the 3D volume to select the slice best representing the center of the tissue, ensuring consistent depth alignment across sessions. Stacks were then cropped to 900×900 pixels in XY, centered on this reference slice, and extracted to a standardized Z-depth of 40 planes for the Day 0 reference timepoint and 80 planes for all subsequent timepoints. Maximum intensity projections (MIPs) were computed along the Z-axis for each channel independently, and per-channel contrast normalization was performed by clipping to the 0.4$^{th}$ and 99.6$^{th}$ intensity percentiles before rescaling to the 0–255 range. A two-channel RGB composite was generated by assigning tdTomato (Channel 1) to red and GFP (Channel 2) to green. Processed stacks were saved as OME-TIFF files with zlib compression and embedded voxel size metadata (0.791 μm/px in XY, 1.6 μm/slice in Z), and composite MIPs were saved as PNG files retaining physical pixel size annotations.

**Cell Count Tracking over Time.** Cell coordinates were identified manually by opening the Day 0 MIP composite images in FIJI and using the multi-point selection tool to place a point at the center of each visible cell in the GFP and tdTomato channels independently. Coordinates were exported as CSV files containing X and Y pixel positions. A binary mask image was then generated from each CSV using a custom Python script employing the scikit-image and imageio libraries, in which a filled circle of fixed radius was drawn at each recorded coordinate on a blank canvas matching the dimensions of the cropped MIP (900×900 pixels). Each circle was assigned a pixel value of 255 against a zero background, producing a binary mask in which each cell was represented by a circular region of interest of 30-pixel radius. Masks were saved as TIFF files and opened as a 20% transparent overlay over the subsequent imaging session MIPs. If cells disappeared under the mask area in subsequent imaging session images, they were counted as a decrease in total cell count for the given channel. Counting was blinded to imaging session, except at the first session, when cell masks were defined.

**Cell Size Tracking over Time.** Changes in cell size over time were quantified using a custom MATLAB script. Maximum intensity projection images from four experimental timepoints (Days 0, 28, 42, 56) were displayed simultaneously in a tiled figure window with linked axes to facilitate direct visual comparison across timepoints. For each cell, a polygon region of interest (ROI) was manually drawn on the Day 0 timepoint image and automatically propagated to the remaining three timepoint images, where it could be independently adjusted to account for any morphological changes or minor spatial shifts. Cell area was calculated from each mask using pixel summation and converted to physical units using the imaging pixel size (0.791 μm/px, yielding 0.626 μm² per pixel). Effective cell diameter was derived from the measured area assuming a circular geometry as $D = 2\sqrt{(A \div \pi)}$. Cell size at each timepoint was additionally expressed relative to the Day 0 measurement to normalize for inter-cell variability in baseline size. All measurements were compiled into a results table and exported as a CSV file containing per-cell area, effective diameter, and normalized relative size across all four timepoints. Soma diameters are relative measures from two-dimensional projections.

***Confocal Images***

Confocal Z-stacks acquired on the Dragonfly 503 were processed as MIPs for each channel prior to analysis. The lesion boundary was delineated manually in FIJI by outlining the GFP-positive region in the green channel, with the resulting outline saved as a binary mask. This mask was used to spatially restrict all subsequent cell counting to the lesion area. A contralateral cortical region was similarly outlined manually and saved as a separate binary mask for use as a within-animal reference.

Cell segmentation was performed using Cellpose (version 3.1.1.2)[10] with custom-trained models for each of the four imaging channels. Separate models were trained from a *cyto3* starting model for the GFP, tdTomato, B-Gal, and pS6 channels to account for the distinct morphological appearances of each cell population. The direct confocal acquisition of B-Gal fluorescence was performed as described in Levitsky *et al.*[11]. Segmentation was run on the single-channel MIPs (*flow_threshold* = 0.5 for GFP and B-Gal; 1.0 for tdTomato and pS6) and the resulting cell outlines were exported as FIJI-compatible ROI ZIP archives for downstream analysis. ROIs with a polygon area below 30 μm², calculated using the shoelace formula and converted to physical units at 0.301 μm/px, were excluded as likely segmentation artifacts. To resolve dual-labeled cells, GFP centroids falling within 5 μm of a tdTomato centroid were reclassified as tdTomato-positive, giving tdTomato identity priority in cases of overlap. For B-Gal, Cellpose ROIs were segmented in FIJI and mean pixel intensity was extracted for each ROI by rasterizing its polygon coordinates onto the single-channel MIP and computing the mean gray value within the resulting binary mask. A fluorescence intensity threshold was then determined by fitting Gaussian distributions separately to the mean gray value distributions of lesion ROIs and contralateral ROIs, with the midpoint between the two distribution means serving as the cutoff. ROIs were additionally required to exceed a minimum area of 30 μm². Only ROIs passing both criteria were retained as B-Gal-positive. An analogous intensity thresholding procedure was applied to pS6, with the contralateral reference derived from raw pixel intensities sampled within the contralateral cortical mask rather than from traced ROIs. NeuN-positive cells were quantified using the *Spots* detection function (Estimated XY Diameter: 10.0 μm; "Quality" filter: 12.0–20.0) in Imaris 10.2 (Oxford Instruments), which identifies cells as discrete puncta based on fluorescence intensity. Spot centroids were exported from Imaris as XLS tables containing physical coordinates, which were converted to pixel coordinates by subtracting the stage offset of the acquired tile and dividing by the pixel size, then filtered to retain only centroids falling within the lesion mask.

Cell densities within the lesion were computed for each marker by dividing the number of centroids inside the lesion mask by the lesion area, calculated by summing mask pixels and converting to $mm^2$. Counts for GFP, tdTomato, and B-Gal were additionally expressed as a percentage of the total NeuN-positive cell counts within the lesion, providing a normalized estimate of each population as a fraction of the total neuronal pool. All results were exported per section as CSV files for statistical analysis.

### ***LC–MS/MS***

Chromatographic data acquisition and peak integration were performed using Analyst 2015 (version 1.6.3; SCIEX). Peaks were integrated automatically with subsequent manual verification. Quantitation was based on analyte-to-internal-standard peak area ratios, with dasatinib-d8 and quercetin-d3 serving as internal standards for dasatinib and quercetin, respectively. Calibration curves were fitted using either linear or quadratic regression with $1/x^2$ weighting, as described above, and final concentrations were back-calculated from the fitted curve parameters within Analyst. No dilution correction factors were applied. Concentrations are reported in nM for both serum and brain matrices.

Samples in which the internal standard signal was absent (serum quercetin, n=9), fell to the lowest percentiles of the batch, or was suspected of a pipetting error (serum dasatinib, n=3) were flagged and excluded from analysis. Samples yielding signals below the lower limit of quantitation were designated as <LOQ and reported alongside the corresponding LOQ value rather than assigned a numerical concentration. Acceptance of each analytical

batch was contingent on satisfactory performance of the quality control samples processed alongside study samples.

#### Statistical Analyses

Statistical tests were performed in MATLAB (version R2024b; MathWorks), Python (version 3.11.5)[12], or GraphPad Prism (version 10.6.1; GraphPad Software, Boston, USA). Histological analyses were performed blind to treatment group (vehicle-only or DQ) and EEG scoring was blind to recording phase; all tests were two-sided. Values are reported as median (IQR) or mean±SEM unless otherwise noted. Nonparametric tests were used throughout, as the small group sizes did not support reliable assessment of normality or equality of variance. Longitudinal cell counts and relative soma diameters were compared across imaging sessions with Friedman tests, and within-animal DQ-phase versus vehicle-phase differences with Wilcoxon signed-rank tests; cell counts in non-FCD controls and NeuN-normalized lesion densities were compared with Kruskal–Wallis tests. Histological cell densities were compared between DQ-treated and vehicle-only animals with Mann–Whitney *U* tests, Bonferroni-corrected across the four markers in Figure 4C and the three markers in Figure 4D. Because lesion size and IUE lesion extent vary between animals, vehicle-only densities were multiplied by a single group-wide factor (1.47) that equated their median tdTomato$^+$ cell density to that of the DQ group before comparison, whereas DQ densities were left unscaled. DQ did not significantly alter tdTomato$^+$ cell number (Figure 2F), so this normalization does not presuppose a treatment effect. LC–MS/MS concentrations were compared across strain and genotype with stratified rank (van Elteren-type) permutation tests, brain and serum values from the same animal with Wilcoxon signed-rank tests, and brain quercetin detection frequency with a stratified exact permutation test. Because each animal contributed a single FOV and baseline cell number differed between fields, group values at each session are reported as means weighted by the number of cells tracked in that animal at the first session ($\Sigma n_i p_i$ / $\Sigma n_i$, where $p_i$ is the percentage of cells remaining in animal *i* and $n_i$ its baseline cell count). Weighting prevents sparsely labeled fields, in which the loss of a single cell produces a disproportionately large percentage change, from dominating the group mean.

Corrections for multiple comparisons were applied where applicable. B-Gal density was the pre-specified primary readout, but Bonferroni correction was applied across all histological markers. *P*-values were taken as significant if less than 0.05 (*), 0.01 (**), and 0.001 (***).
